# Automated decoding the mouse body postures in a whisker nuisance test

**DOI:** 10.1101/2022.11.27.518077

**Authors:** Gabriele Chelini, Tommaso Fortunato-Asquini, Ottavia Ollari, Tommaso Pecchia, Yuri Bozzi

**Affiliations:** Center for Mind/Brain Sciences - CIMeC, University of Trento, Corso Bettini 31, 38068 Rovereto (TN), Italy; Department of Cellular, Computational, and Integrative Biology (CIBIO), University of Trento, via Sommarive 9, 38123 Trento, Italy; CNR Neuroscience Institute, via Moruzzi 1, 56124 Pisa, Italy

**Keywords:** autism, somatosensory, whisker response, behavioral analysis, deeplabcut

## Abstract

Abnormal response to sensory stimuli characterizes multiple neuropsychiatric conditions. However, not many tools are currently available to assess somatosensory abnormalities in rodent models of brain disorders, limiting the possibilities to study cellular and molecular correlates of this phenotypic trait. To this goal, previous studies relied on the whisker nuisance test (WNt), in which freely moving mice are constantly stimulated on their whiskers using a wood stick for a set time. The whisker-guided response is then deconstructed in behavioral categories indicative of either anxiety or curiosity. Thus far, WNt was shown to be a valuable tool to investigate sensory-driven abnormalities in mouse models of autism spectrum disorders (ASD), demonstrating a solid translational validity. Nevertheless, assessment of behavioral response in the WNt is currently limited by the lack of an objective quantification method. To overcome this limitation, we developed WNt3R (Whisker Nuisance Test in 3D for Rodents), a MATLAB toolbox that uses the output of the open-source software DeepLabCut to determine discrete body postures associated with multiple ethologically-relevant behaviors. Our results show that behavioral modules identified using WNt3R reliably decode mouse body postures, outperforming the manual user, thus offering a novel and unbiased approach to study altered somatosensory function in rodents.

## Introduction

The use of animal models to study brain disorders often struggles with the need for translational validity, limiting the investigator’s ability of extrapolating valuable mechanistic information about the neural correlates of pathological behaviors. Using a very simple approach, the whisker nuisance test (WNt), previous studies identified a variety of whisker-dependent phenotypes in mouse models of autism spectrum disorders (ASD)^1–3^. This test consists in manually stimulating the mouse whiskers with a wood stick over 3 consecutive, 5-minutes long sessions, while the animal is freely moving in a delimited space. Mouse response to the whisker stimulation is then quantified using pre-determined behavioral categories (BCs). BCs used thus far aim to deconstruct the animals’ response by exploring independent instances of either anxious or explorative behaviors^1–4^. Previous studies have shown how behavioral abnormalities identified with the WNt are reliably well-matched with similar observation in people with ASD^5–8^, showing a solid translational validity and opening the possibility of directly investigating the neurobiological underpinning of this common phenotypic trait^1–3^. Nevertheless, thus far, behavioral quantification of WNt exclusively relied on subjective judgment assigning a numerical value to multiple behavioral manifestations^1–4^. To this goal, an experimenter was trained on recognizing and deciphering mouse postures with reliable and systematic confidence. This approach, other than being massively time-consuming, comes with considerable limitations such as inter- and intra-individual variability, introduction of personal biases and is forcingly limited by the number of pre-determined behavioral categories.

To overcome these constrains, we conceptualized an automated procedure to interpret mouse postures on a sub-second scale, using a combination of novel machine-learning based behavioral tracking and unsupervised cluster analysis. After visualizing approximately 100 videos, from as many animals, we concluded that one of the best predictors of mice behavior during WNt could be simplified as the spatial location, along the vertical (z)-axis, of specific ‘bodily hotspots’: nose (No), eyes (E), neck (N), middle of the back (mB), lower back (lB), base of the tail (T) (Fig.1a-c). More specifically, the relative location of one hotspot compared to the others changes drastically according to specific behavioral states such as spatial navigation, explorative rearing or passive avoidance (Fig.1a-c). Taking advantage of the new-generation software for pose estimation *DeepLabCut* (DLC)^9,10^, we tracked these hotspots during the whisker stimulation and used the output coordinates to dynamically read mice body language during the test. Here we describe the results of such analysis, showing that behavioral modules automatically identified using WNt3R can reliably decode mouse body postures.

**Fig.1.**
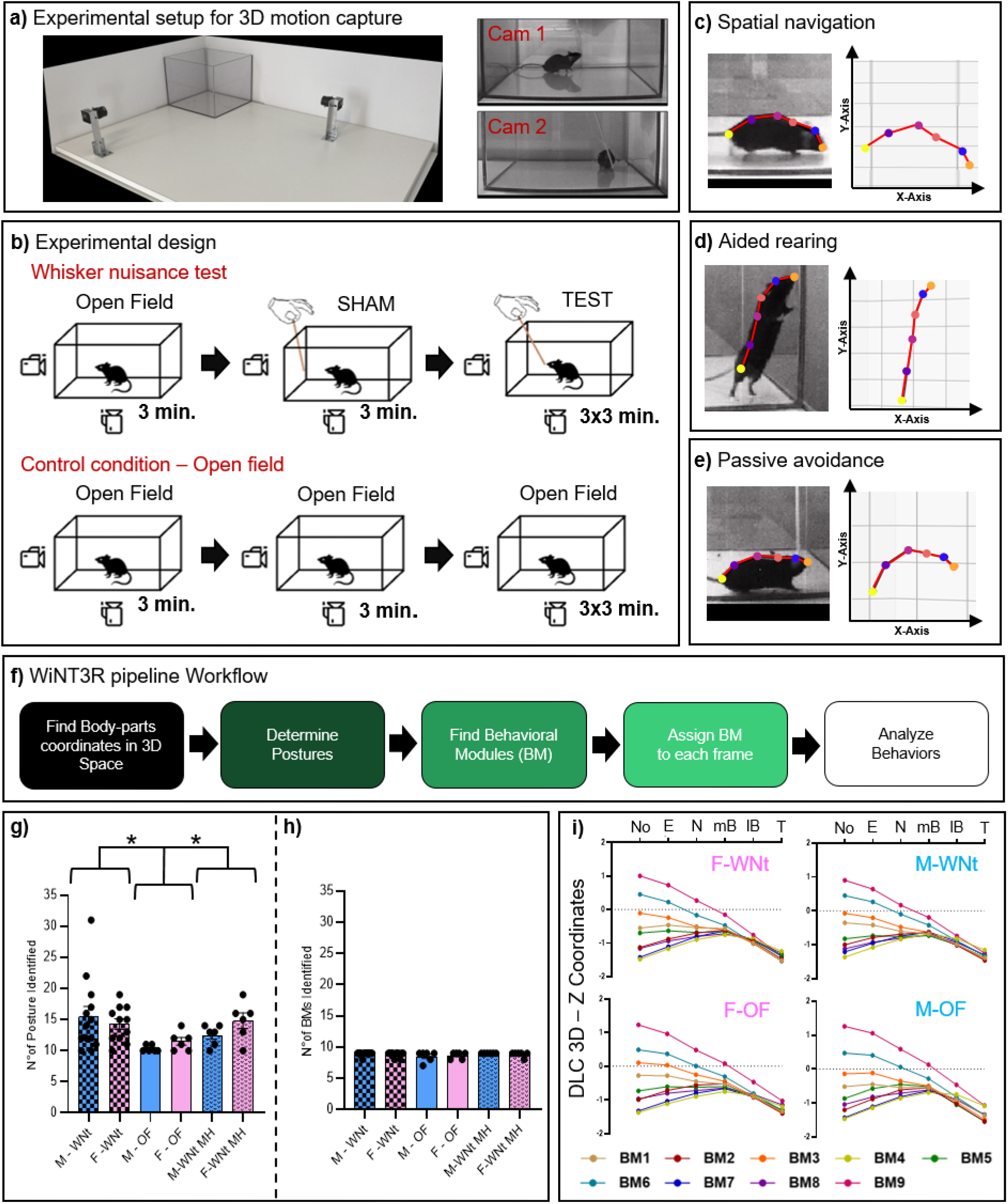
Methodological apparatus of the WNt3R approach. **a)** Angular view of the WNt3R arena (left). Note the stereo-configuration of the cameras for 3D-motion capture. Orthogonal view of the same mouse during the WNt (right). **b)** Experimental design **c-e)** Example of body-posture determined by visual inspection (left) and corresponding coordinates detected by DLC-3D. Bodily-hotspot are indicated by colored dots (Orange=Nose; Blu=Eyes; Pale-Red=Neck; Magenta=Mid-back; Purple=Lower-back; Yellow=Base of the tail). **f)** Schematic workflow of the WNt3R pipeline. **g-i)** A two-step cluster analysis eliminates inter-individual variability **(g)** and identifies replicable and comparable BMs **(h) i)** Schematic view of BMs identified during whisker stimulation and control conditions, for both sexes.

## Materials and Methods

### Animals

Animal research protocols were reviewed and approved by the University of Trento animal care committee and Italian Ministry of Health, in accordance with the Italian law (DL 26/2014, EU 63/2010). Animals were housed in a 12h light/dark cycle with unrestricted access to food and water. All sacrifices for brain explant were performed under anesthesia and all efforts were made to minimize suffering. A total of 48 age-matched adult wild-type littermates of both sexes (weight 25–35 g) were used for the study. All mice were generated from our CNTNAP2 colony (C57BL/6 background)^11^. Thirty-six mice were assigned to the whisker stimulation test, while 12 did not receive any stimulation and served as controls.

### Whisker nuisance test (WNt)

WNt was performed with some variation compared to what previously described^1,2^. Animals were daily habituated to the experimental room for a week prior the test. Starting four days before the test, mice were daily habituated to the experimental cage (a fiberglass cubic box with open top; 25×25×20 cm) and the experimenter for 2 minutes. This time was extended to 10 minutes the day before the test. This habituation scheme will be referred as heavy-habituation (HH) for the rest of the manuscript. On test day, mice were acclimated with the experimental environment for 10 minutes, before the beginning of the testing phase. Before the beginning of the actual test, three minutes of the animal freely moving in the open field (OF) arena were videotaped as a baseline activity session. The testing phase was composed of four sessions, lasting 3 minutes each. In the first session (SHAM), a wood stick was introduced in the experimental cage, avoiding direct contact with the animal. The following three sessions consisted in stimulating mice’s whiskers by continuously deflecting vibrissae using the wooden stick (at a frequency of approximately 3 strokes per second). An additional group of mice was run in the same experimental conditions, but without any exposure to the wood stick. This OF variant of the test was used as a control condition to assess the specificity of the whisker-guided response compared to spontaneous freely-moving activity. Finally, to assess the efficacy of WNt3R in detecting anxiety-related behaviors, an additional group of 12 mice underwent the WNt after only 2 days of habituation (mild habituation – MH).

To dissect the complexity of behavioral response, multiple behavioral categories were identified and independently quantified by a trained observer. The identified categories, based on a previous version of the test^1–4^ included fearful and curious behaviors. Fearful behaviors were divided *in_active avoidance* (time the animal spends actively avoiding the stick) and *passive avoidance* (time the animal spends in a defensive posture consisting in curved back, protracted neck and stretched limbs or retracted in fully hunched posture; this measure also included the time in freezing). Curious behaviors were divided in *aided rearing* (when the animals, during a rearing action, leans on the arena’s walls investigating the surrounding environment) and *unaided rearing* (when the rearing is not supported by walls; this behavior usually occurs toward the center of the arena and is often associated with stick exploration during the WNt).

### 3-D motion capture in the WNt3R arena

A basic requirement, for Deeplabcut 3D (DLC-3D) tracking, is to capture the subject motion using (at least) two videocameras in a stereo-configuration^9,10^. To this goal, two videocameras (ImageSource DMK27BUR0135 USB3-monochrome, equipped with TAMRON 13VM308ASIRII objectives) were secured, using a metal pedestal, on a board of laminated wood, in orthogonal positioning respect to each-other (Fig.1a). At the corner of the base, two aluminum guides were placed to ensure the stability of the cubic cage during the test. This allowed us to replace the experimental cage at every given animal, thus eliminating experimental biases due to the odor of other mice, that could potentially alter the emotional state of the test subject. Video were acquired using the OBS-studio software and edited using OpenShot Video Editor, before being analyzed with DLC. For details about how to use DLC-3D we recommend using this link: https://github.com/DeepLabCut/DeepLabCut/blob/master/docs/Overviewof3D.md

### Tissue processing and immunofluorescence

Ninety minutes after the end of the WNt, mice were deeply anesthetized with isoflurane and sacrificed by decapitation. Brains were excised, washed in 0.1% phosphate buffer saline (PBS) and post-fixed overnight in 4% paraformaldehyde (PFA), switched to a cryoprotectant solution (80% PBS, 20% glycerol with 0.1% sodium azide) and stored at 4°C. Cryoprotected brains were sectioned on a vibratome (Leica, VT1200) at 40μm thickness. Serial sections were collected in 24 separate compartments and stored at 4°C in cryoprotectant solution.

Free-floating slices were rinsed three times in PBS (10 min each), then washed in PBS containing 0.2% detergent (Triton-X, Fisher, AC215680010)) for 30 minutes. Tissue sections were then incubated in blocking solution [2% Bovine serum albumin (BSA), 1% fetal bovine serum (FBS) in PBS] for 3 hr and then transferred to primary antibody solution [2% BSA, 1% FBS, 1 to 1000 dilution of primary antibody (Rabbit anti-Arc/Arg 3.1, Proteintech - Cat.16290-1-AP)] and incubated at room temperature for 24 hr. Then, sections were rinsed three times in PBS (5 min each) and placed in a fluorophore-conjugated (Alexa Fluor^TM^ Plus 488) secondary antibody solution [1:300 dilution of donkey anti-Rabbit secondary antibody (Thermo Fisher, AB_2762833) in PBS] for 24 hr. Sections were then washed 5 min in PB, mounted on superfrost slides, dried for 1 hr and coverslipped with fluorescent mounting medium (Southern biotech 0100-01). Slides were stored at 4°C in the dark until used.

### Confocal microscopy and image analysis

A confocal laser scanning microscope Leica TCS-SP8, equipped with a HC PL APO 63x/1.40 oil objective and interfaced with Leica “LAS-X” software was used to detect ARC immunolabeling in the Lateral and Basolateral nuclei of the amygdala. Images were recorded at a resolution of 1024 pixels square, 400 Hz scan speed. Excitation/emission wavelengths were: 490/520 for Alexa-488 fluorophore. Acquisition parameters were set during the first acquisition and kept consistent for all the images. Corrected fluorescence intensity (CFI) was quantified using imageJ software according to the formula: CFI = integrated density – (area size*mean fluorescence of background readings).

### Data Collection and Analysis

Data collection for all studies was carried out by investigators blind to experimental conditions. All statistical analyses were carried out using Prism8 software (GraphPad Software San Diego, CA)

## Results

### WNt3R identifies meaningful behavioral modules (BMs) repeated across animals

The final goal of the method we are proposing is to analyze the time spent by the animals in specific body postures, in order to compare the identified behavioral categories across experimental groups. This was achieved using a two-step cluster analysis (Fig.1f). Using the raw DLC-3D files, WNt3R pipeline generates a matrix containing the linear distance between each hotspot with every other. Using this distance matrix, an unsupervised k-means clustering^12^ identifies a *k* number of postures using the elbow method, a statistical strategy, originally proposed by Robert Thorndike, to determine the most appropriate number of *k-mean* clusters to choose^13^. This within-animal analysis generates a certain degree of inter-individual variability, limiting the capacity to compare behaviors across experimental groups (Fig.1g). To solve this problem, a second clustering is run ‘between-animals’, taking as an input the averaged distance matrix for each identified posture, for any given animal. With this step, individual postures are matched and allocated into behavioral modules (BMs) replicated across all (or most) animals, allowing for comparative analysis (Fig.1h). Interestingly, our data show that the number of identified postures significantly differ between experimental condition (WNt or OF), demonstrating the WNt3R efficacy to identify specific stimulus-driven behaviors (Fig.1g).

### WNt3R automatically discriminates between experimental conditions

To assess WNt3R ability to discriminate between experimental conditions, we compared the percentage of time spent in each BM during the habituation phase with the four consecutive sessions of the test. Predictably, no significant differences were observed at any time-point in the OF condition (Fig.2-left; p-values are summarized in Fig. S1-left). Conversely, we identified marked and significant changes in the HH (BM1,3,5,7,8,9) and MH (BM6,8,9) groups at several different time-points (Fig.2-middle and right; all p-values for this analysis are summarized in Fig.S1). Interestingly, the majority of the BMs (7 out of 9) show significant changes starting from the SHAM session, when the stimulus (stick) is introduced in the arena, but before the beginning of the whisker stimulation. This data demonstrates that the exposure to a novel stimulus, although not tactile, is sufficient to trigger a meaningful change in mice behavioral state.

**Fig.2.**
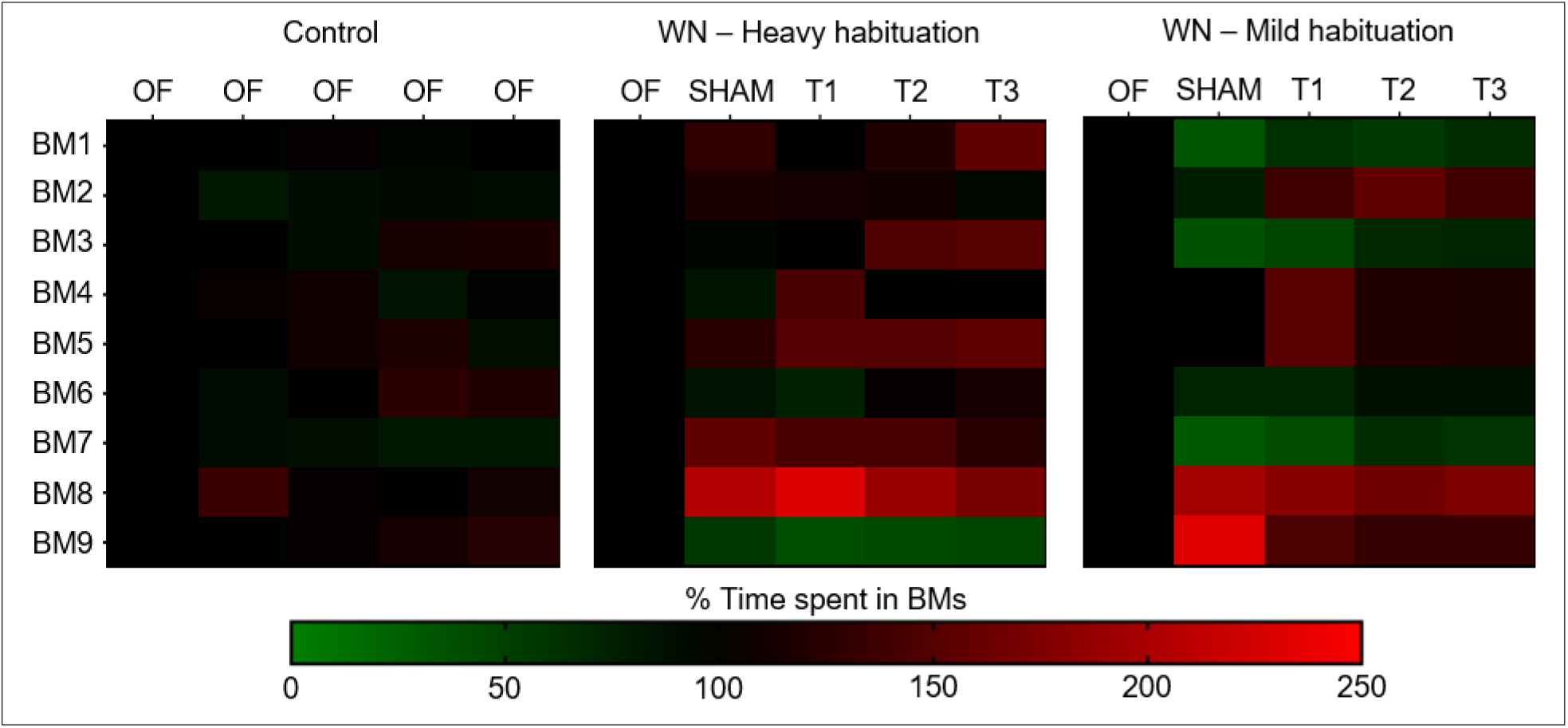
Time spent in each BM changes drastically depending on the test session and experimental conditions.

### Isolating the whisker-guided response results in time-dependent changes in BMs

To extrapolate the whisker-guided component to the behavioral changes, we expressed BMs intensity as variation in frequency (Δf) normalized over their frequency (f) during the SHAM session. As a reference baseline we established that there was no significant change in the control condition (open field with no whisker stimulation). In the MH, instead, marked significant changes were found for BM1 (T1-T2-T3), BM3 (T2-T3), BM5 (T2-T3) and BM8 (T2-T3), compared to SHAM. Finally, in HH, differences with SHAM were observed in BM1 (T3), BM3 (T2-T3), BM4 (T1), BM5 (T1-T2-T3) and BM6 (T3). Altogether these findings confirm that WNt3R classification method successfully discriminates time- and stimulus-driven changes in the expression of specific behaviors. (All data are summarized in Fig.3 a-i. p-values are summarized in Fig.3j and Table S2)

**Fig.3.**
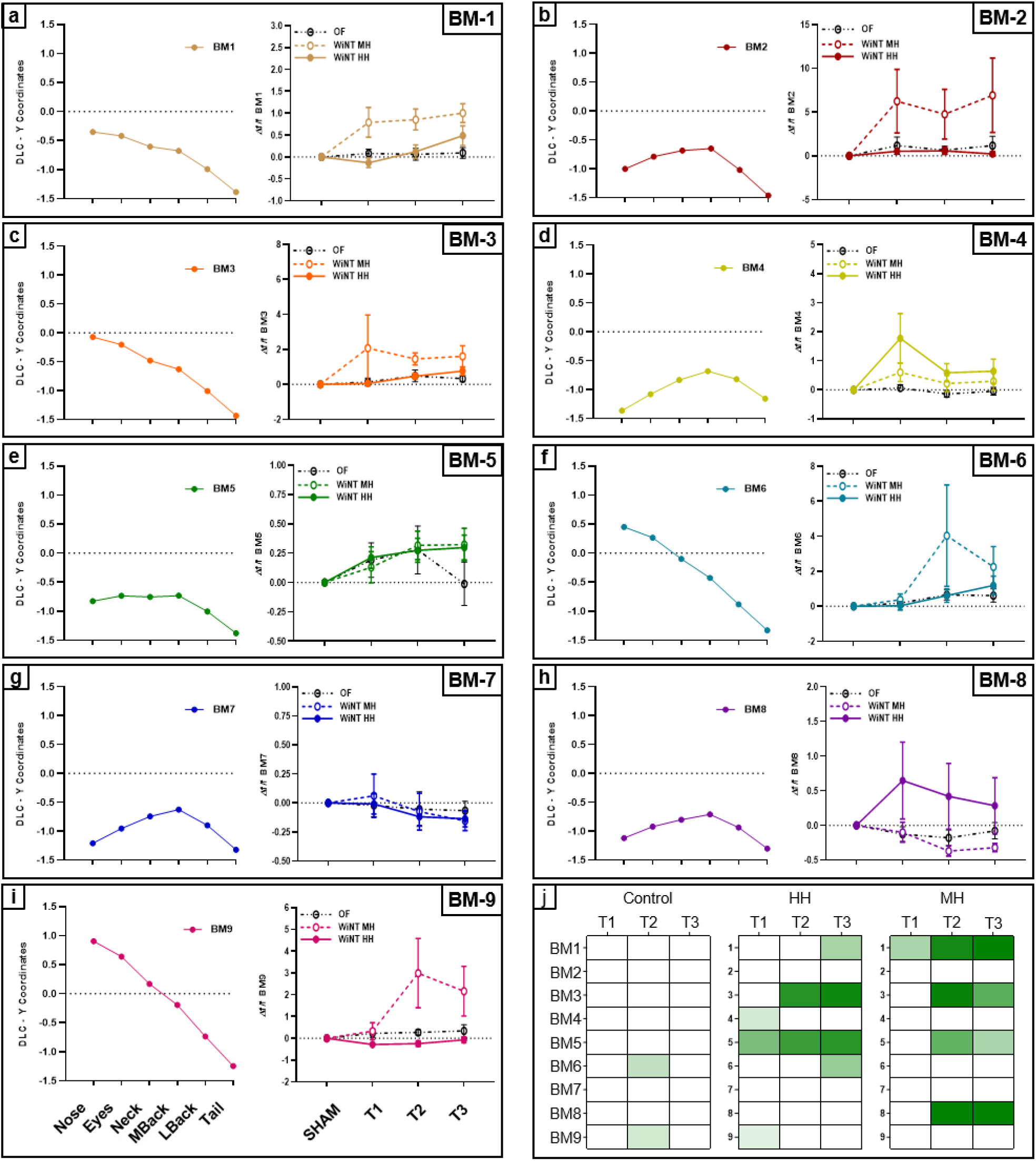
Isolation and analysis of the whisker-guided response. For each panel **a-i):** the left plot shows a schematic view of body-parts coordinates plotted in 2D. The right plot shows the Δf/f of BM with respect to the SHAM session, over the 3 consecutive trials of the WNt. **i)** Heatmaps showing significant differences obtained comparing the Δf/f of each trial with the baseline value (SHAM) (darker green shows values further from p=0,05).

### BMs are informative of mice emotional response to the whisker stimulation

Aiming to use WNt3R to identify dynamic changes in the mouse emotional state, we sought to determine whether specific BMs could associate with discrete BCs quantified by a manual user. To do so, we generated a matrix correlating the outcome of the manual quantification for each BC with each independent BM (Fig.4a). We observed that BCs associated with fear and anxiety had a strong positive correlation with 2 out of the 9 BMs (4-8), while being negatively correlated with categories 1-3-6-9 (Fig.4a and Table S2). More specifically, among fear-related BMs, BM4 showed stronger correlation with evasive behaviors (r=0.42, p=0.0002. Fig.4b) while BM8 showed stronger association with passive avoidance response (r=0.36, p=0.001. Fig.4c). A mirrored trend was observed for explorative BCs (Fig.4a and Table S3), where strong positive correlations were found for BMs 1-3-6-9 (Fig.4d shows correlation between BM6 and total rearing time, r= 0.92, p<0.0001, while negative correlations was observed for BMs 2-7-8. Only one BM (5) showed no correlation with either BCs, most likely indicative of an emotionally neutral behavior such as spontaneous spatial navigation. (p-values are summarized in table S3). These data confirm that BMs identified by WNt3R can be used as a proxy to isolate multiple ethologically-relevant behaviors, allowing comparative studies.

**Fig.4.**
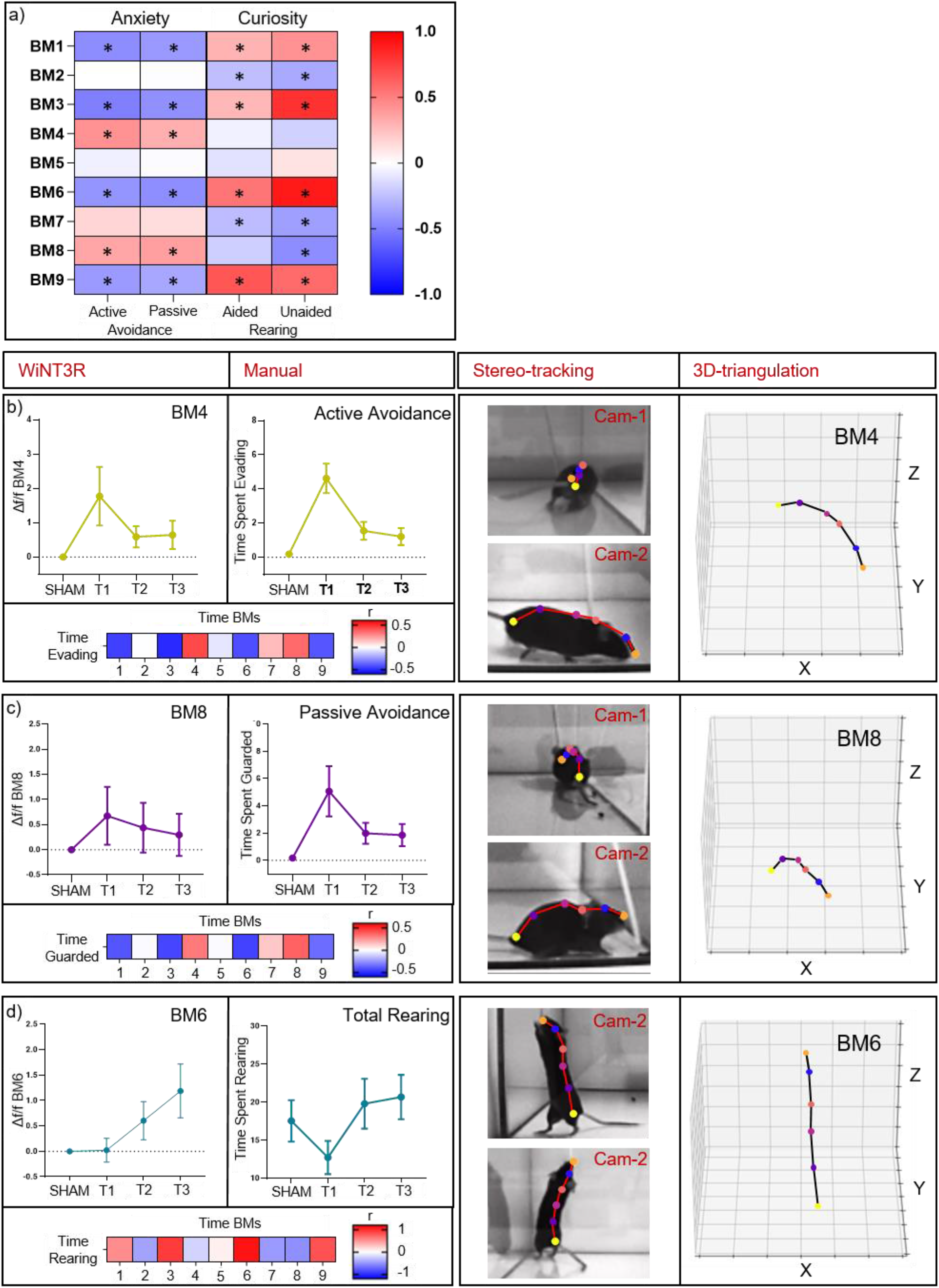
BMs correlate with discrete BCs quantified by a trained user. **a)** Heatmap showing correlation between BMs and BCs. Asterisk indicates significance of the correlation. Color indicates strength and direction of Pearson-r. **b) c) d)** **Left panel – Top charts**: correlated BMs and BCs show a similar temporal progression **Left panel – Bottom chart:** heatmaps showing the strength and direction of the correlations between BMs and BCs **Right panel:** representative frames corresponding to each BMs displayed in the left panel.

### Fear-associated BM8 finds direct neural correspondence in the basolateral amygdala

Finally, as a proof-of-principle of the biological relevance of our behavioral classification, we asked whether BMs could be mapped within specific brain areas. To do so we assessed the association between the activation of the amygdaloid complex with freezing-associated BM8 (Fig.4a,c), aiming to find correspondence with anxiety-related behaviors. To probe for neural activation, we used immunofluorescent labeling of ARC, a neuroplastic protein involved in stress-response in the amygdala^14^, consistently with previous studies showing increased expression of immediate-early genes in response to fear^15^ and whisker stimulation^2^. We quantified the fluorescent intensity within the lateral (Lat) and basolateral (BLA) nuclei. Strikingly, in BLA, we found that ARC-expression was selectively positively correlated with BM8 (r=0.81, p=0.001. Fig.5a,b). To the contrary, BLA-ARC expression was negatively correlated with BM6 (r=-0,61, p=0,05. Fig.5a), which was identified as indicative of explorative behavior. No significant correlation was found between Lat-ARC expression and neither of the BMs (data not shown). These findings suggests that blindly-identified BMs correspond to biologically-relevant instances represented within specific neural circuits.

**Fig.5.**
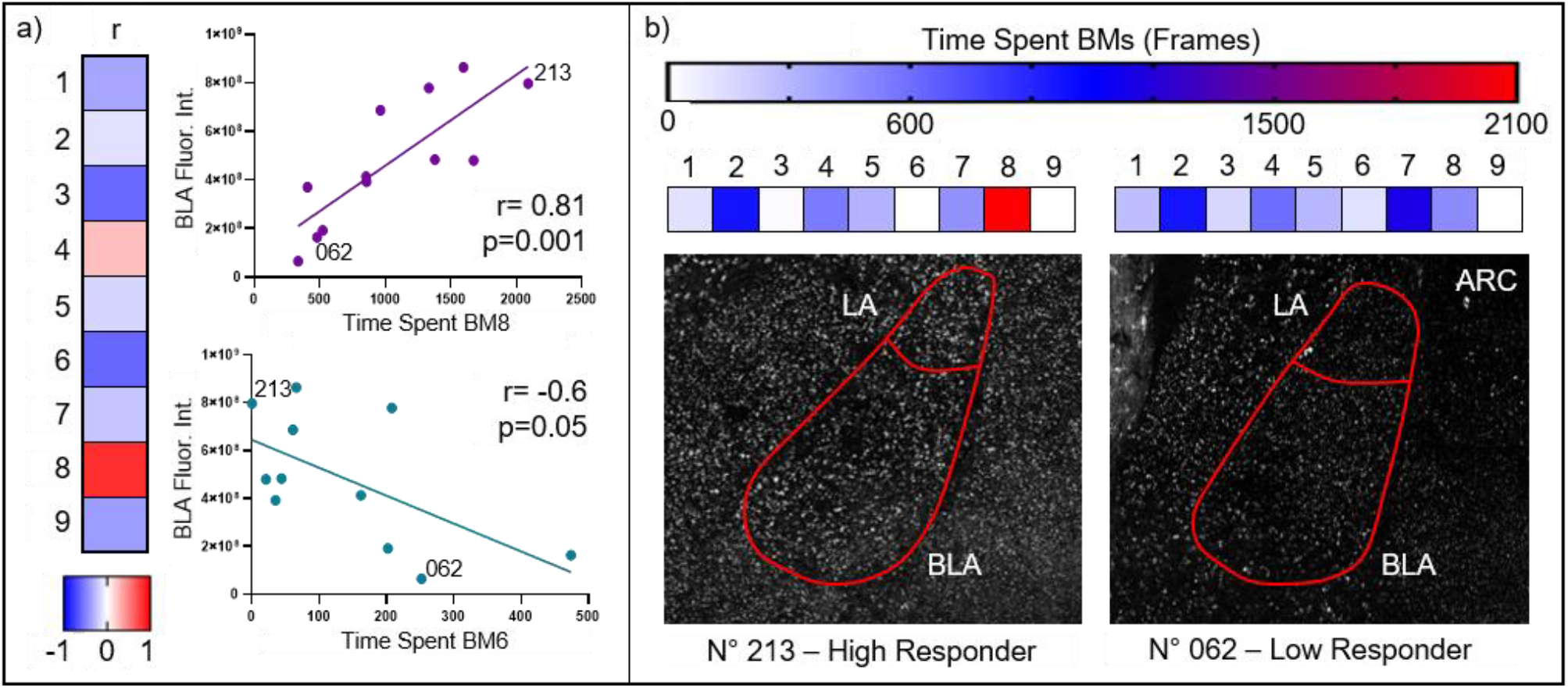
**a)** Arc protein expression in BLA correlates positively with BM8 and negatively with BM6. **b)** Representative picture depicting one animal displaying elevated fearful response and BLA activation (left, N°213) compared to a second subject showing low fearful response and low BLA activation (right, N°062).

## Discussion

In this work we used a new-generation software for mouse body-tracking combined with a strategy for automated posture detection to achieve successful and biologically relevant deconstruction of the mouse body-language during a simple whisker-dependent test. This approach comes with numerous advantages. First, automated behavioral detection eliminates any individual biases by determining replicable and meaningful categories while significantly reducing the time required for quantification. Moreover, a major limitation of the manual quantification consists in the fact that most BCs show virtually the same temporal progression; anxiety-related behaviors present dramatic increase during the first trial followed by a sharp decline in the second trial due to adaptation^**1,2**^; an opposite tendency is observed for explorative categories^1^. The unbiased extraction of BMs depicts behavioral categories characterized by their own independent temporal fluctuation as well as specific intrinsic inter-individual variability, demonstrating a more scrupulous selection of behavioral states. At the same time, the automated quantification retains the capacity of differentiating between experimental groups, as shown by the drastic changes observed between HH and MH (Fig. 2). With this piece of evidence, we additionally confirmed the necessity of a careful and drastic habituation phase to eliminate non-specific anxious behaviors potentially masking whisker-guided anxiety (Fig. S1). Moreover, we were able to identify marginal, but significant sex-differences in two of the BMs (6-9, Fig.S2), despite a relatively small sample size, unveiling an important piece of evidence missing from previous studies. Given parallelism and discrepancies observed with the manual quantification, it is important to clarify the divergent nature of BMs with respect to traditional behavioral characterization. Especially for anxiety-related behaviors we have found significant correlation with BCs, but the strength of the correlation remained weak (less than 0.5 Fig. 4a-c). This is due not only to the unavoidable noise included in both measures, but mostly to the fact that the two measures are not directly overlapping. BMs are, by definition, indirect measures that can be used as a proxy to infer emotional states, they are not the behavioral state itself. By the same principle, correlation between manually or automatically quantified rearing is virtually perfect, as in that case the visual observer is directly quantifying the posture rather than a more complex behavioral construct.

Further developments of WNt3R will aim to increase the complexity of the posture descriptors to achieve stronger deconstruction of mouse body-language. As an important confirmation of the biological relevance of BMs, we showed that the frequency of BM8, putatively identified as associated with passive avoidance response, was strongly correlated with activity-dependent ARC-immunolabeling in BLA. Our results are in agreement with previous studies showing that the expression of immediate-early genes, including ARC, is induced in somatosensory areas^16–19^ and amygdala^2^ in response to whisker stimulation.

While further studies are needed to determine whether similar brain-behavioral correlation are detectable for each individual BM, the strength of this association works as a proof-of-principle showing that BMs are not only valuable proxies for behavioral interpretation, but have a well-defined neuroanatomical location that can be exploited to investigate the biological underpinning of abnormal whisker-guided response. Furthermore, this observation opens the door for future studies aimed to explore how the neural activity in BLA affect the dynamic transition between behavioral states during whisker stimulation.

The biggest limitation of WNt3R consists in its intrinsic reductionist approach. Previous studies demonstrated how mice behavior can be decomposed in several meaningful sub-second structures^20^. Conversely, WNt3R reduces this complexity in an array of interpretable behavioral modules. While limited in the ability of interpreting mice behavior, we showed how this approach allows the user to achieve a meaningful interpretation of behavioral flow, to be used for translational inference. On the technical level, a limitation of WNt3R is given by a certain degree of error due to the fact that some data points (frames) are forcingly compressed within behavioral categories. This results in datapoints contributing to expand the noise contained within singular behavioral modules. This sets an aim for future development of WNt3R: by discretizing the current classification it would be theoretically possible to include additional principal components to the cluster analysis improving the precision of behavioral detection. An additional weakness of our approach is the nature of the stimulation itself. To provide continuous whisker stimulation in freely-moving rodents we are currently bound to manually deliver the stimulation, losing the possibility of controlling for important factors such as the number of whisker deflections, intensity and frequency of the stimulus presentation, and continuity of the stimulation overtime. Therefore, the WNt is not suitable for systematic investigation of the whisker sensation and its neural correlates. It has been, however, previously shown the capacity of this test to identify multiple sensory-driven endophenotypes in rodent models of ASDs^1–4^. Finally, our study specifically focused on the whisker-guided response, which is not, *per-se*, a salient emotional stimulus and only triggers dramatic avoidance response in a subset of wild-type mice. However, given our success in detecting emotional-relevant BMs in a mildly-emotionally charged task, we have reason to believe that the same approach could be expanded to other stimulus-driven behaviors. Changing the modality of stimulus presentation (such as auditory or olfactory) should not affect the detection of emotional states; on the contrary, the presentation of emotionally salient stimuli might result in a more distinctive behavioral parcellation. More generally, the method here proposed might allow to improve the translational validity of a number of behavioral tests by reducing quantification biases, ensure replicability (within and beyond different laboratories) and expand the number of behavioral categories to be analyzed.

## Supporting information

Clip BM4

Clip BM9

Clip BM6

## Funding

GC effort is covered by ‘CARITRO postdoctoral fellowship’, funded by Fondazione Cassa di Risparmio di Trento e Rovereto. YB is supported by the Strategic Project TRAIN - Trentino Autism Initiative (https://projects.unitn.it/train/index.html) from the University of Trento (grant 2018-2022).

## Acknowledgments

We thank all the administrative and technical staff of CIMeC for support. A special thank goes to Mrs. Michela Maffei, animal caretaker of the CIMeC animal facility, for her endless care and support provided in managing the animal colony used in the study. We thank carpenter Mr. Marco Chelini for donating materials and equipment used in the study.

## Supplementary Material

**Fig. S1.**
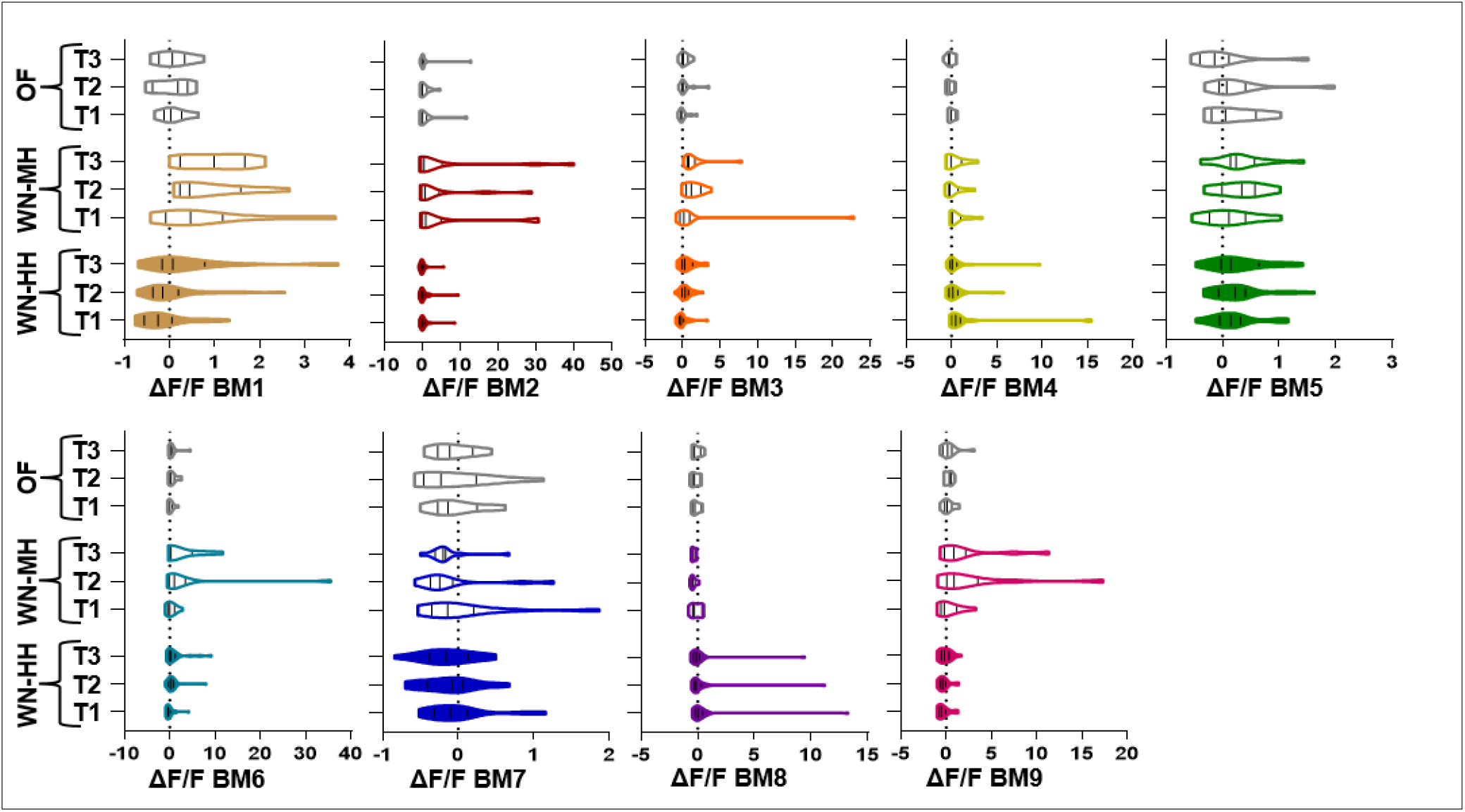
Violin plots depicting the dispersion of the data across experimental conditions. Notably, anxiety-related modules (BM4 and BM8) show no data spread in the MH condition, suggesting that the anxiogenic effect of the whisker stimulation is masked by excessive generalized anxiety expressed during the SHAM session.

**Fig.S2.**
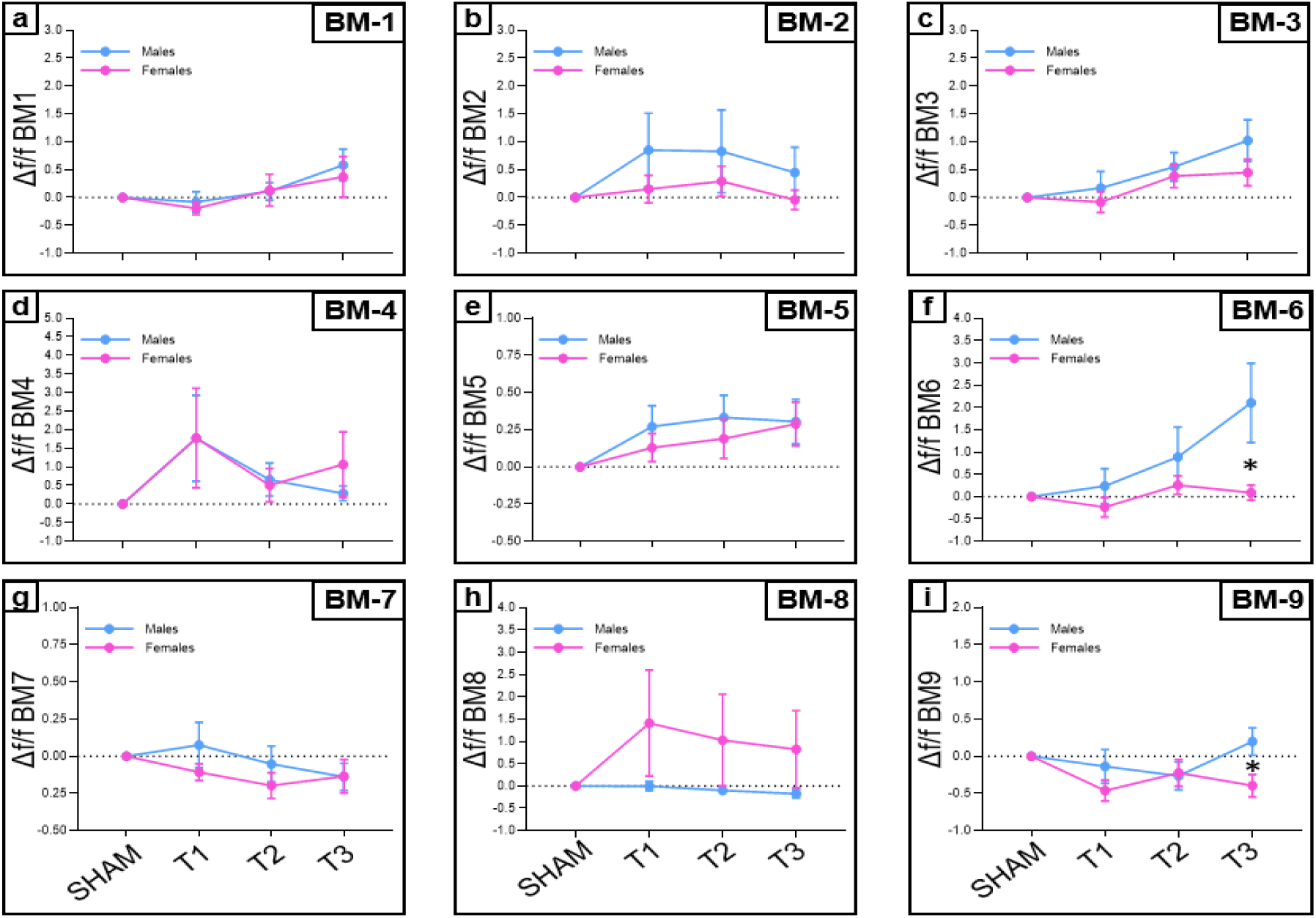
Sex Differences. Only marginal significant differences can be observed between sexes (BM6-T3 and BM9-T3). * Indicates p-values lower than 0.05.

**Table S1.** p-values obtained comparing the initial habituation session (OF) with each test sessions for every BM across all 3 experimental conditions. Bold indicates p-values lower than 0,05.

|  | Control |  |  |  | WN – Heavy Habituation |  |  |  | WN – Mild Habituation |  |  |  |
| --- | --- | --- | --- | --- | --- | --- | --- | --- | --- | --- | --- | --- |
|  | OF | OF | OF | OF | SHAM | T1 | T2 | T3 | SHAM | T1 | T2 | T3 |
| BM1 | 0,65 | 0,47 | 0,85 | 0,49 | 0,10 | 0,93 | 0,16 | **0,01** | 0,52 | 0,50 | 0,27 | 0,11 |
| BM2 | 0,46 | 0,44 | 0,389 | 0,39 | 0,40 | 0,34 | 0,47 | 0,68 | 0,66 | 0,11 | 0,10 | 0,15 |
| BM3 | 0,85 | 0,96 | 0,12 | 0,21 | 0,74 | 0,94 | 0,06 | **0,01** | 0,17 | 0,30 | 0,30 | 0,34 |
| BM4 | 0,30 | 0,32 | 0,55 | 0,63 | 0,20 | 0,05 | 0,92 | 0,91 | 0,28 | 0,08 | 0,51 | 0,29 |
| BM5 | 0,86 | 0,42 | 0,28 | 0,08 | 0,05 | **0,03** | **0,00** | **0,02** | 0,07 | 0,12 | 0,93 | 0,91 |
| BM6 | 0,68 | 0,67 | 0,07 | 0,09 | 0,28 | 0,06 | 0,81 | 0,52 | **<0,01** | **<0,01** | 0,89 | 0,15 |
| BM7 | 0,61 | 0,94 | 0,66 | 0,86 | **0,01** | **0,03** | 0,14 | 0,15 | 0,10 | 0,16 | 0,44 | 0,37 |
| BM8 | 0,07 | 0,28 | 0,58 | 0,17 | **<0,01** | **<0,01** | **<0,01** | **<0,01** | **<0,01** | **0,00** | **0,02** | **0,02** |
| BM9 | 0,4 | 0,21 | 0,09 | 0,18 | **<0,01** | **<0,01** | **<0,01** | **<0,01** | **<0,01** | **<0,01** | 0,02 | 0,04 |

**Table S2.** p-values obtained comparing the Δf/f of each session with the baseline value 0 obtained in the SHAM session. Each table refers to a different experimental condition across all experimental conditions (data are summarized in fig,2j).

|  | Control |  |  | WN – Heavy Habituation |  |  | WN – Mild Habituation |  |  |
| --- | --- | --- | --- | --- | --- | --- | --- | --- | --- |
|  | OF | OF | OF | T1 | T2 | T3 | T1 | T2 | T3 |
| BM1 | 0,422 | 0,655 | 0,467 | 0,242 | 0,439 | **0,039** | **0,040** | **0,004** | **0,001** |
| BM2 | 0,261 | 0,165 | 0,297 | 0,171 | 0,174 | 0,396 | 0,117 | 0,124 | 0,135 |
| BM3 | 0,566 | 0,185 | 0,099 | 0,767 | **0,009** | **0,003** | 0,297 | **0,001** | **0,021** |
| BM4 | 0,583 | 0,252 | 0,735 | **0,049** | 0,069 | 0,133 | 0,085 | 0,445 | 0,367 |
| BM5 | 0,225 | 0,206 | 0,953 | **0,029** | **0,014** | **0,01** | 0,35 | **0,023** | **0,04** |
| BM6 | 0,366 | **0,045** | 0,135 | 0,925 | 0,123 | **0,036** | 0,268 | 0,192 | 0,079 |
| BM7 | 0,837 | 0,730 | 0,431 | 0,927 | 0,134 | 0,064 | 0,748 | 0,65 | 0,091 |
| BM8 | 0,258 | 0,141 | 0,526 | 0,245 | 0,366 | 0,473 | 0,51 | **0,001** | **<0,001** |
| BM9 | 0,237 | **0,049** | 0,261 | 0,053 | 0,066 | 0,59 | 0,411 | 0,089 | 0,087 |

**Table S3.**
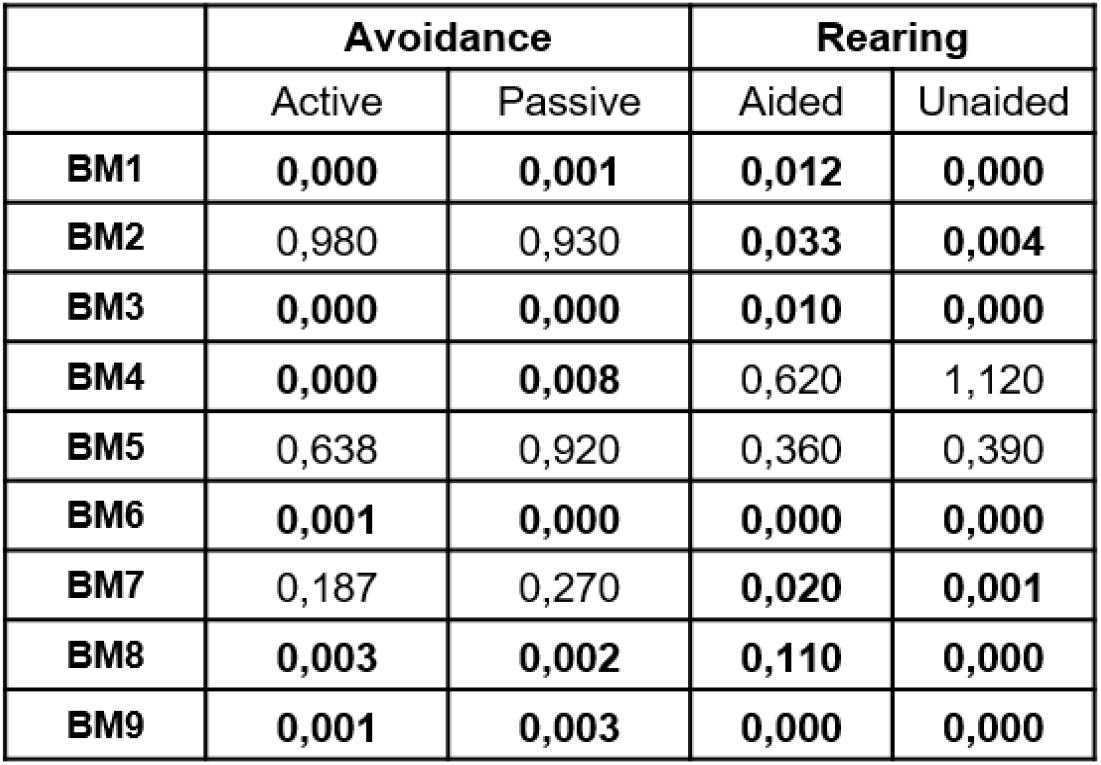
p-values of the correlation between BMs and BCs. (Bold indicates values lower than 0.05)

|  | Avoidance |  | Rearing |  |
| --- | --- | --- | --- | --- |
|  | Active | Passive | Aided | Unaided |
| BM1 | **0,000** | **0,001** | **0,012** | **0,000** |
| BM2 | 0,980 | 0,930 | **0,033** | **0,004** |
| BM3 | **0,000** | **0,000** | **0,010** | **0,000** |
| BM4 | **0,000** | **0,008** | 0,620 | 1,120 |
| BM5 | 0,638 | 0,920 | 0,360 | 0,390 |
| BM6 | **0,001** | **0,000** | **0,000** | **0,000** |
| BM7 | 0,187 | 0,270 | **0,020** | **0,001** |
| BM8 | **0,003** | **0,002** | **0,110** | **0,000** |
| BM9 | **0,001** | **0,003** | **0,000** | **0,000** |

## Notes

**Competing Interest Statement:** The authors declare no competing interests.

### Competing Interest Statement

The authors have declared no competing interest.

